# Event-related potential (ERP) evidence of early visual processing differences in cystinosis

**DOI:** 10.1101/2023.03.31.535154

**Authors:** Douwe J. Horsthuis, Sophie Molholm, John J. Foxe, Ana A. Francisco

## Abstract

Cystinosis, a rare lysosomal storage disease, is characterized by cystine crystallization and accumulation within tissues and organs, including the kidneys and brain. Its impact on neural function appears mild relative to its effects on other organs, but therapeutic advances have led to substantially increased life expectancy, necessitating deeper understanding of its impact on neurocognitive function.

Behavioral difficulties have been reported in cystinosis in the visual and visual-processing domain. Very little is known, however, about how the brains of people living with cystinosis process visual information, although cysteamine accumulation in the retina is a prominent feature of cystinosis. Here, electrophysiology was recorded during a Go/No-Go task to investigate early visual processing in cystinosis, compared to an age-matched control group. Analyses focused on early stages of cortical visual processing.

The groups differed in their initial cortical response, with individuals with cystinosis exhibiting a significantly larger visual evoked potential (VEP) in the 130 to 150 ms time window. The timing and topography of this response suggested an enhanced P1 in cystinosis that could be the result of cortical hyperexcitability and/or differences in attentional engagement and explain, at least partially, the visual and visual-spatial difficulties described in this population. The groups also differed in the associations between neural responses and verbal abilities: While controls with higher IQ scores presented larger neural responses, that relationship was not observed in cystinosis.

## INTRODUCTION

Cystinosis is an autosomal recessive disorder caused by bi-allelic mutations in the 17p13.2-located CTNS gene (1) with an incidence rate of around one in 100,000 to 200,000 live births (2, 3). The accumulation of cystine in cells (4, 5)—characteristic of the condition—leads to deregulation of endocytosis and cell signaling processing (6). Ultimately, intralysosomal cystine crystallizes, triggering significant damage in a multitude of tissues and organs (7). Renal, retinal, endocrinological, muscular, and neurological complications are observed in cystinosis (8, 9). Due to drug development advances and the availability of renal replacement therapy (10, 11), the life expectancy of individuals with cystinosis has, however, increased well into adulthood, demanding a better understanding of the developmental trajectories associated with the condition.

We have been particularly interested in characterizing the impact of cystinosis in brain and cognitive function. Though abnormally high levels of cystine have been observed in different brain regions (12–14), cystinosis’ impact on brain activity is still not well understood. In our previous work, mainly focused on auditory processing and sensory memory, we reported maintained sensory processing, but some differences in sensory memory in children and adults living with cystinosis (15, 16).

Here, we expand this investigation to the visual sensory domain. Interestingly, a differential between visual-perceptual and verbal indices has been often reported in this population, with the former being significantly lower than the latter (17–19). This pattern appears to emerge early in development and to persist throughout the lifespan (20, 21), regardless of age at treatment onset (22). Significant difficulties have also been described in visual-motor, visual-spatial and visual memory skills (22–26), but see (27) for a report of maintained visual learning in adults with cystinosis. Despite this pattern of relative weaknesses in visual processing domains, very little is known about how the brains of people living with cystinosis process visual information. One case study tested visual processing in two children with cystinosis before and after kidney transplantation. During dialysis treatment, both children showed delayed and decreased early visual evoked responses, but typical brain responses were seen after kidney transplantation (28). Despite the very small number of individuals tested to date, EEG measurements appear to be sensitive to neuropathology in cystinosis and could be useful as outcome measures to assess the impact of treatment on brain function.

Here, we use high-density EEG to investigate basic visual processing in a group of individuals with cystinosis and compare them to a group of age-matched controls. The analyses focus on the visual evoked potentials (VEPs) P1 and N1. P1 is an early VEP peaking around 100 ms following stimulus onset and has been associated with multiple generators in both dorsal and ventral visual streams (29–32). N1 is the negative-going component that follows P1 and peaks at around 160 ms after stimulus onset. It seems to be primarily generated in structures in the ventral visual stream (30), as evidenced by intracranial grid-electrode recordings (33) and scalp topographic studies (34, 35). Additionally, we tested for associations between neural responses and age and standardized cognitive measures.

Considering the behavioral difficulties reported in visual-related processing in cystinosis and the tendency for cysteamine accumulation in the retina, we hypothesized that individuals with cystinosis would show different early sensory-perceptual brain responses to visually presented stimuli, reflected in amplitude differences in the VEPs P1 and N1. Identifying the processing stages that are impaired and contribute to visual and visual-spatial processing differences in cystinosis is crucial to develop impactful strategies to address the difficulties reported in previous behavioral studies and to identify sensitive biomarkers of treatment efficacy on brain function.

## MATERIALS AND METHODS

### Participants

Thirty-eight individuals diagnosed with cystinosis (CYS; age range: 7-36 years old, 25 women) and 45 age-matched controls (CT; 27 women) were recruited. Individuals with cystinosis were recruited via social media and through contact with family organizations. Due to the rareness of cystinosis, most participants, all of whom lived within the continental United States, traveled from out-of-state to participate. Furthermore, the controls were recruited via flyers in the neighborhoods surrounding the lab and through a lab-maintained participant database. Developmental and/or educational difficulties or delays, neurological problems, and a severe mental illness diagnosis were exclusionary criteria for controls. Current neurological problems were exclusionary criteria for individuals living with cystinosis. To assess visual acuity, a Snellen chart was used. All participants had normal or corrected to normal vision. Nevertheless, all participants were asked at the start of the EEG paradigm if they could see the stimuli and their different components without difficulty. One individual with cystinosis was excluded from the final sample due to illness on the scheduled day of testing. All individuals, and their legal guardian if under 18 years old, signed a consent form. Participants were monetarily compensated for their time. This study and all the associated procedures were approved by the Albert Einstein College of Medicine Institutional Review Board. All aspects of the research conformed to the tenets of the Declaration of Helsinki.

### Experimental Procedure and Stimuli

Participation consisted of two visits, which involved completion of a cognitive function battery and EEG recordings. The cognitive function battery included the assessment of verbal and non-verbal intelligence (using age-appropriate Wechsler Intelligence Scales) and executive functioning components (Delis-Kaplan Executive Function System, D-KEFS; (36) and the Conners Continuous Performance Test 3, CPT; (37)) (to be reported elsewhere). During the EEG recording session, participants were asked to respond to different tasks assessing sensory processing and response inhibition. Here, we focus on basic visual processing of images presented in the context of a Go/No-Go task. Response inhibition findings are reported separately.

During the EEG Go/No-Go task, positive and neutral valence images from the International Affective Picture System (IAPS; Lang and Cuthbert, 1997) were presented in a pseudorandom sequence. Participants were instructed to press the left mouse button upon each stimulus presentation, as quickly and as accurately as possible, unless the stimulus was a repetition of the immediately preceding stimulus, in which case they should inhibit their response. Stimuli, subtended 8.6° horizontally by 6.5° vertically, were presented centrally every 1000 ms for 600 ms with a (random) inter-stimulus-interval between 350 and 450 ms. Three 12-minute blocks were run. Each block consisted of 540 trials, for a total of 1620 per participant.

Here, we focus exclusively on hit trials, that is, trials that included a correct button press after stimulus presentation, to assure that participants were paying attention and were engaged in the task. Additionally, hits were defined as responses to a non-repeated picture, when preceded by another hit to guarantee that those responses were not impacted by inhibitory processes from a previous trial.

### Data acquisition and analysis

Continuous EEG data were recorded from 64 scalp electrodes at a sampling rate of 512 Hz (Active 2 system; Biosemi™, The Netherlands; 10-20 montage) and preprocessed using the EEGLAB toolbox (version 2021.0) (38) for MATLAB (version 2021a; MathWorks, Natick, MA) (the full pipeline can be accessed at: https://github.com/DouweHorsthuis/EEG_to_ERP_pipeline_stats_R) (39). Preprocessing steps included down-sampling data to 256 Hz, re-referencing to the average, and filtering with a 0.1 Hz high pass filter (0.1 Hz transition bandwidth, filter order 16896) and a 45 Hz low pass filter (11 Hz transition bandwidth, filter order 152). Both were zero-phase Hamming windowed sinc FIR filters. Noisy channels were excluded based on kurtosis and visual confirmation. Artifacts from blinks and saccades were eliminated via Independent Component Analysis (ICA). The spherical spline method was then used to interpolate channels that were removed in previous steps. Data were segmented into epochs of −100 ms to 400 ms using a baseline of −100 ms to 0 ms. These epochs went through an artifact detection algorithm (moving window peak-to-peak threshold at 120 μV). To equate number of trials per participant, 200 trials for hits were chosen randomly per subject.

P1 was measured between 130 and 150 ms and N1 between 165 and 190 ms at O1, Oz, and O2. Time windows and electrode locations were selected based on past research and confirmed (and adjusted) by inspecting the timing and topography of the major voltage fluctuations in the grand averages. Mean amplitude data were used for both between-groups statistics and Spearman correlations. All *p-values* (from *t*-tests and Spearman correlations) were submitted to Holm-Bonferroni corrections for multiple comparisons (40), using the *p.adjust* of the *stats* package in R (41). Mixed-effects models were implemented to analyze trial-by-trial data, using the *lmer* function in the *lme4* package (42) in R (41). Group was always a fixed factor. Subjects and trials were added as random factors. Models were fit using the maximum likelihood criterion. *P* values were estimated using *Satterthwaite* approximations.

## RESULTS

### Demographics and cognitive function measures

Table 1 shows a summary of the included participants’ age, sex, and cognitive functioning (verbal and non-verbal IQ). Two-sample independent-means *t* tests were run in R (41) to test for group differences in age and cognitive performance. In cases in which the assumption of the homogeneity of variances was violated, *Welch* corrections were applied to adjust the degrees of freedom. A chi-square test was run to test for independence between sex and group. Effect sizes were calculated utilizing Cohen’s *d* and *w*. As can be seen in Table 1 and Figure 1, the groups differed in verbal and non-verbal abilities, with individuals with cystinosis showing more difficulties than their age-matched peers. No differences were found in age and sex between the groups.

**Figure 1.**
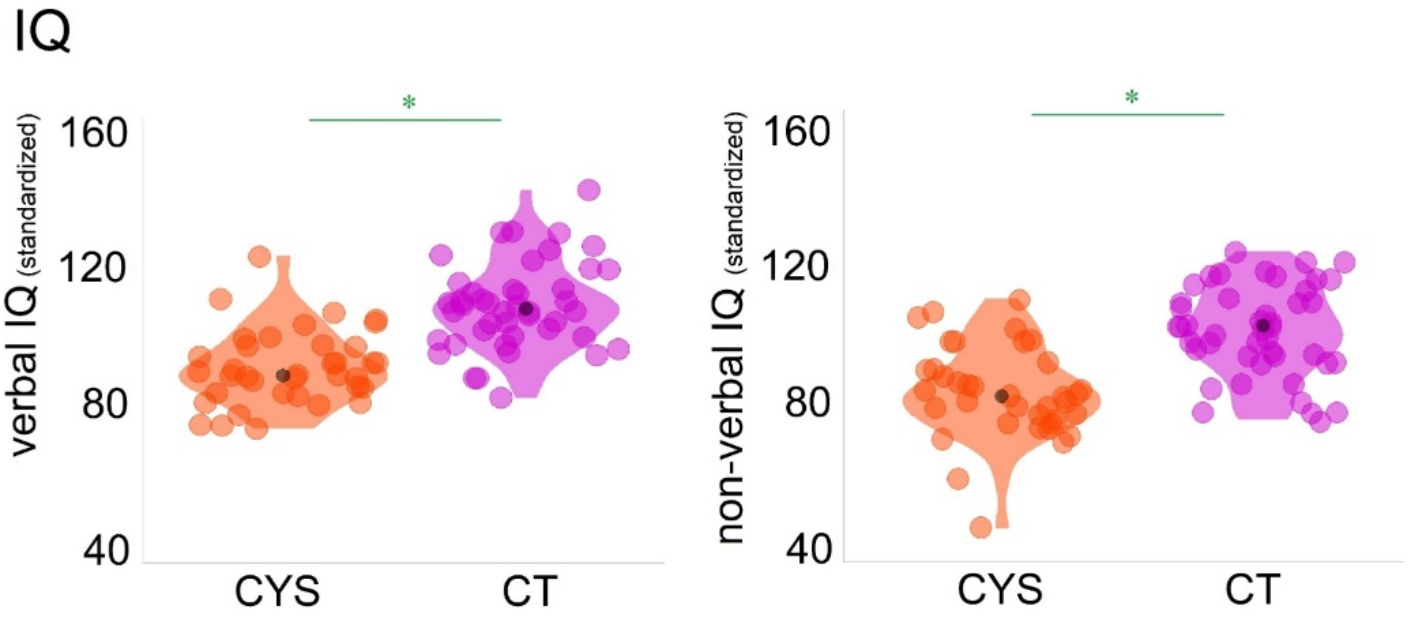
Included participants’ verbal and non-verbal IQ scores.

**Table 1.** Characterization of the control and cystinosis individuals included in the analyses: Demographics and cognitive function (IQ and inhibition measures).

|  | control | cystinosis | statistical test | effect sizes |
| --- | --- | --- | --- | --- |
| **age** | M=17.36; SD=8.92 | M=17.62; SD=9.36 | $t=-0.13, df=75.43, p=.90$ | $d=0.03$ |
| **sex** | 27 F, 18 M | 25 F, 12 M | $\chi^2=0.23, df=1, p=.63$ | $w=0.08$ |
| **verbal IQ** | M=109.71; SD=19.85 | M=93.46; SD=11.21 | $t=4.66, df=71.57, p<.01$ | $d=1.00$ |
| **non-verbal IQ** | M=103.44; SD=13.49 | M=85.19; SD=13.70 | $t=6.04, df=76.50, p<.01$ | $d=1.36$ |

### Basic visual processing: P1 and N1

The averaged ERPs for P1 and N1 per channel and by group can be seen in Figure 2. Mixed-effects models were implemented as described in the Methods Section. Average amplitude at O1-Oz-O2 at each trial was the numeric dependent variable. As can be appreciated in the waveforms and scalp topographic maps in Figure 2, individuals with cystinosis showed significantly increased P1 amplitudes compared to their age-matched peers (*β* = 7.55, SE = 2.45, *p* = .01). A peak-to-peak measure (subtracting the amplitude of the N1 from the amplitude of the P1) was made for the N1, to control for amplitude differences in the P1. After controlling for the differences seen in P1, the groups did not significantly differ in the N1 time window (*β* = −1.41, SE = 0.90, *p* = .12).

**Figure 2.**
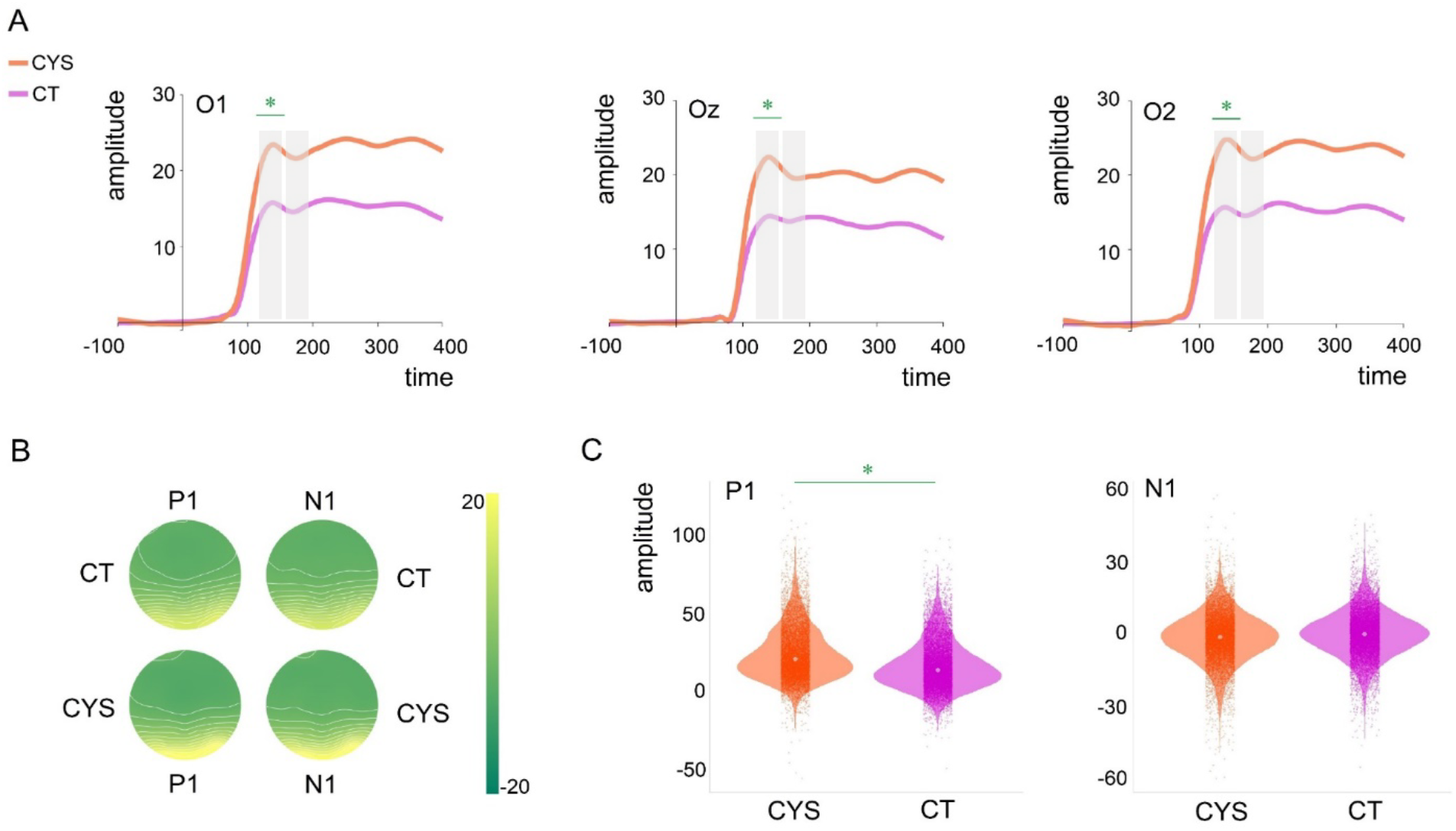
Panel A: Averaged ERPs per group and channel (O1, Oz, and O2). Shaded areas indicate windows of interest. Asterisks indicate significant differences. Panel B: Topographical maps showing brain activity in the P1 and N1 time windows per group. Panel C: Violin plots showing distribution of single trial data amplitudes per group at O1, Oz, and O2 (average) for P1 and N1. Small dots indicate amplitude at a given trial and central dots group mean amplitude. Asterisks indicate significant differences.

#### Correlations

Figure 3 shows neural responses (P1 and N1) correlations with age (Panel A), verbal IQ (Panel B), and non-verbal IQ (Panel C). P1 and N1 amplitudes significantly correlated with age in both groups: While P1 decreased with age (*r_s_*=−.73, *p* = .01), N1 increased with age (*r_s_*=.40, *p* = .02). As can be seen in Figure 3 Panel B, the groups differed in the correlations between P1/N1 and verbal IQ. While controls showed a positive correlation between P1 and verbal IQ (*r_s_*=.40, *p* = .04) and a negative correlation between N1 and verbal IQ (*r_s_*=−.45, *p* = .02), such correlations were not significant in cystinosis. Neither P1 nor N1 correlated significantly with non-verbal IQ. This pattern was observed in both control and cystinosis groups.

**Figure 3.**
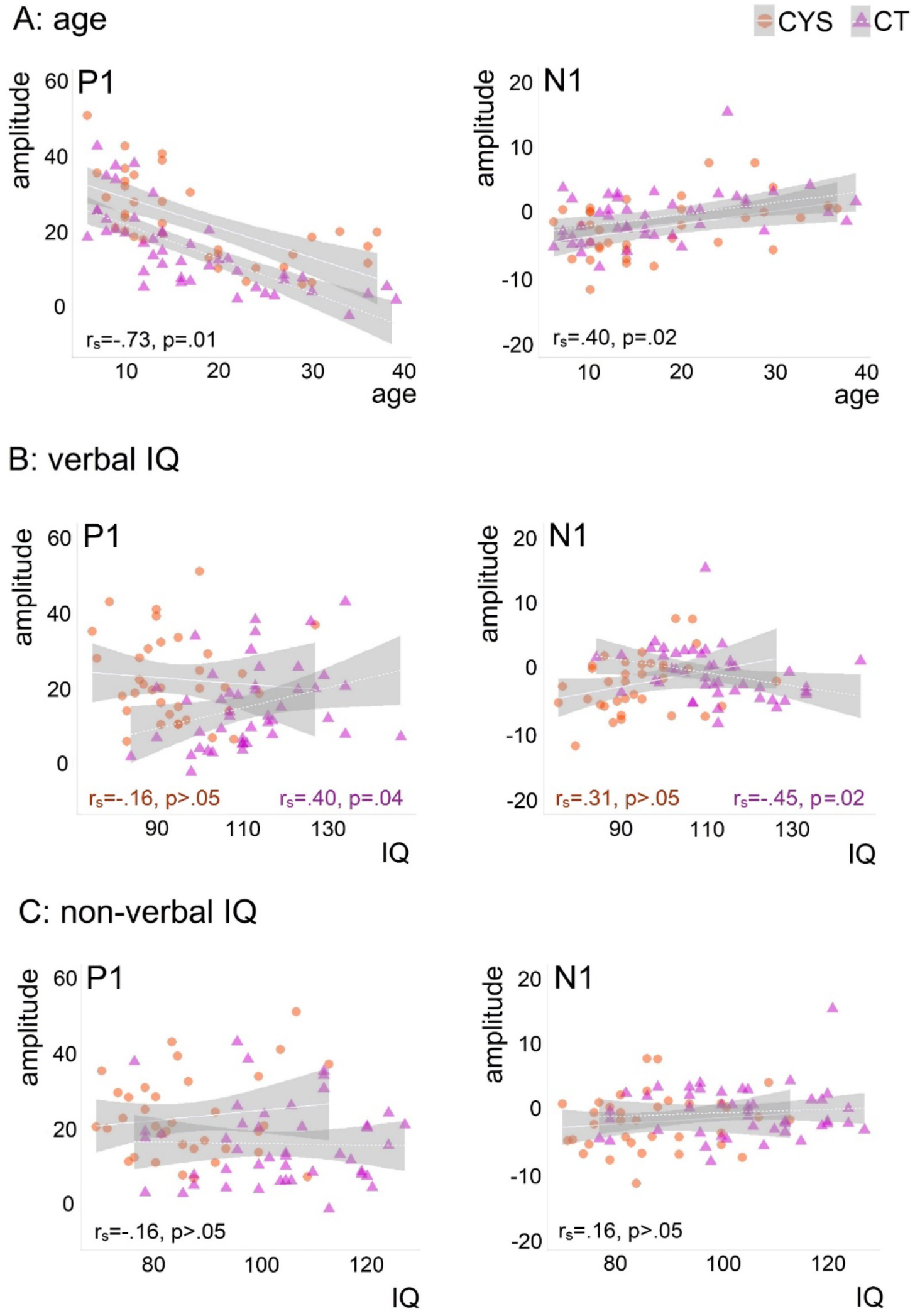
Spearman correlations between P1 and N1 and age (Panel A), verbal IQ (Panel B), and non-verbal IQ (Panel C). Correlation coefficients and their significance are presented for the full sample when groups did not differ (age and non-verbal IQ) and per group when groups differed (verbal IQ).

### DISCUSSION

To better characterize previously reported visual-spatial difficulties in cystinosis (20, 26, 43), we investigated visual processing in this population by focusing on early electrophysiological brain responses, the P1 and N1, to visually presented stimuli.

Differences in the VEP in the P1 time window were found between those with and without cystinosis, with individuals with cystinosis showing significantly increased P1 amplitudes. Increased early sensory-evoked potentials may reflect increased excitability in sensory cortices (44–48). An increased P1 in cystinosis could thus signify visual cortex hyperexcitability in this population. Visual cortex hyperexcitability has been linked to photophobia in individuals suffering from migraines (44), who may also present with increased visual evoked potentials (such as the P1)—though not consistently so (see (45) for a summary of different results). Importantly, photophobia is the most frequently reported ocular symptom in cystinosis (3, 49, 50). Though we have not objectively quantified photophobia in our cystinosis sample, comments regarding general sensitivity to light were common during testing. The pathophysiology of photophobia in relation to crystal accumulation in cystinosis is unclear. Nevertheless, it appears that, in this population, enhanced early visual processing could be related to hypersensitivity to light and/or visual cortex hyperexcitability. Future EEG research with this population should take sensitivity to light into consideration when designing visual experiments. Additionally, these findings and their implications in classroom and professional settings utilizing digital media and screens should be further explored.

Differences between the groups in P1 amplitudes could additionally reflect different levels of attentional engagement. Attention can significantly modulate P1 amplitudes (51–54). Such modulation may reflect a sensory gain type mechanism that results in enhanced perceptual processing of attended stimuli (55). Interestingly, we have previously reported increased P2 and P3a amplitudes in a sample of adults with cystinosis who performed an auditory duration oddball task (15). We argued then that individuals with cystinosis may engage attention differently, which the current findings could be further evidence of. However, one might expect such modulation to be likewise present during the N1 time window (e.g., (56)). Studies focused on attentional processes in this population are needed, particularly those that relate to potential differences in attention engagement with the cognitive profile associated with cystinosis.

P1 has been localized to sources in the dorsal extrastriate cortex, specifically to areas V3, V3a, and adjacent middle occipital gyrus and in the ventral extrastriate cortex, specifically area V4 in the fusiform gyrus (29–32) and it is thus driven by both magnocellular and parvocellular input (57). Traditionally, V3 and V3a have been associated with motion processing (e.g., (58). But studies in non-human primates suggest that these areas may also be involved in linking higher-level parietal and temporal processing streams (59) and may play a role in the integration of visual stimulus features, essential for global stimulus processing (60). V4 has been associated with the processing of surface properties (color, brightness, texture), shape (orientation, curvature), motion and motion contrast, and depth; but it has also been shown to play a crucial role in visual attention (61). Moreover, in non-human primates, V4 appears to be widely interconnected with other visual areas along the ventral and dorsal visual streams, frontal areas, and subcortical structures, and thus has been conceived as holding an integrative role in visual perception and recognition and, potentially, in guiding perceptual decisions and higher-order behavior (62). Structural and/or functional neuroimaging studies examining V3, V3a, and V4 could be particularly informative in understanding the mechanistic roles that these areas play in the visual-spatial and visual attention differences described in cystinosis.

In the N1 time window, after accounting for amplitude differences between the groups in the earlier VEP, no differences were found between individuals with cystinosis and their age-matched control peers. Accounting for those earlier differences was fundamental given that, as can be appreciated in Figure 2A, differences between the groups emerge around the onset of P1 and remain stable for the remainder of the analyzed epoch. If one considers in isolation the numeric difference between the two groups at distinct time points after P1, these would be significant. However, we believe that differences that onset around the P1 time window dictate these later differences in amplitude. Future work should be directed at further characterizing visual processing differences and their implications for higher order processes such as object localization and identification.

Of note, this pattern of increased amplitudes in cystinosis appears to be especially localized to the occipital and lateral parietal-occipital channels. While the current analyses focused on the channels in which response was maximal for both groups, those channels also represented the largest differences between the groups. As one can appreciate in Figure S1 (supplementary materials), differences between the groups are reduced in lateral parietal-occipital channels and absent in mid-line parietal-occipital and across parietal channels. Future studies directed at defining the underlying neural sources explaining these differences and lack thereof are justified.

Correlational analyses revealed different associations between age, neural responses, and clinical measures. An important finding is that neural and cognitive function correlated with age in a similar fashion across groups. In the general population, VEPs appear to remain stable until early adulthood, at which time their amplitudes begin to attenuate (63). Here, such attenuation was also observed in the cystinosis group. Likewise, there was no evidence of cognitive decline with age in cystinosis, at least not one that deviates from what is expected in the general population. The groups differed, however, in the correlations between P1/N1 and verbal IQ. While controls with higher IQ scores showed larger P1s and N1s, such relationship was not present in the group of individuals with cystinosis. As seen in Figure 3B, in cystinosis, although not statistically significant, increased N1 could be indicative of cognitive function differences. This finding speaks to the presence of different mechanisms at play in the relationship between cognitive abilities and early visual processing in cystinosis.

Lastly, a brief mention to findings relating to general cognitive function in cystinosis. Neurocognitive assessments showed lower verbal and non-verbal IQs in individuals with cystinosis, when compared to their control peers. We and others had previously reported lower IQ scores in this population (15, 16, 19, 23) and, as in other studies, our findings (see Table 1) indicate greater difficulties in nonverbal processing (18–22, 27). Of note, scales as the Wechsler Scales of Intelligence have an unbalanced number of timed non-verbal vs verbal subtests. To more accurately characterize the cognitive profile associated with cystinosis, it would be important to investigate whether, given the time, individuals with cystinosis would still show marked difficulties in non-verbal tasks or whether such difficulties are mainly explained by processing speed differences. While this would complicate comparisons with normative data, it could be useful to understand discrepancies between verbal versus non-verbal subtests.

This study is not without limitations. First, our groups were not matched in terms of IQ, with the cystinosis group presenting significantly lower scores than the control group. Such differences could have impacted the results. Second, variables related to current health status (such as a measure of renal function) and compliance to treatment, which has been linked to better clinical outcomes (64), were not included in the present study but could be useful in understanding group- and individual-level differences. Lastly, most of the individuals with cystinosis, and differently from those included in the control group, travelled to the lab from out-of-state and completed the study in two days in a row, which could have increased tiredness in this group. Regardless, here, we present seminal evidence of early visual processing differences in cystinosis which could contribute to, at least partially, the behavioral visual-spatial difficulties described in this population. More work is needed to describe how very early visual processing differences might contribute to the cognitive profile associated with cystinosis. A better understanding of such associations could contribute to the identification of sensitive biomarkers of treatment efficacy on brain function. The current findings should motivate future studies utilizing paradigms tapping into higher- and lower-order visual processes and investigating visual processing and pathways in more depth, for instance, by employing tasks distinguishing between ventral and dorsal streams and magnocellular and parvocellular pathways

## ACKNOWLEDGEMENTS

We wish to thank Dr. Juliana Bates, who performed the clinical assessments, Elise Taverna, Danielle Newbury, and Alaina Berruti for their help with data collection and Dr. Frederick J. Kaskel for his help with recruitment. We extend our most sincere gratitude to the participants and their families for their interest, their involvement, and their time. This work was supported by a grant from the Cystinosis Research Network and a Eunice Kennedy Shriver National Institute of Child Health and Human Development U54 Grant (HD090260) to the Human Clinical Phenotyping Core of the Rose F. Kennedy Intellectual and Developmental Disabilities Research Center.

## COMPETING INTERESTS

The authors declare no conflicts of interest.

## AUTHOR CONTRIBUTIONS

AAF, SM, and JJF conceived the study. DJH and AAF collected and analyzed the data. DJH and AAF wrote the first draft of the manuscript. AAF, SM, and JJF provided editorial input to DJH on the subsequent drafts.

## SUPPLEMENTARY MATERIALS

**Figure S1.**
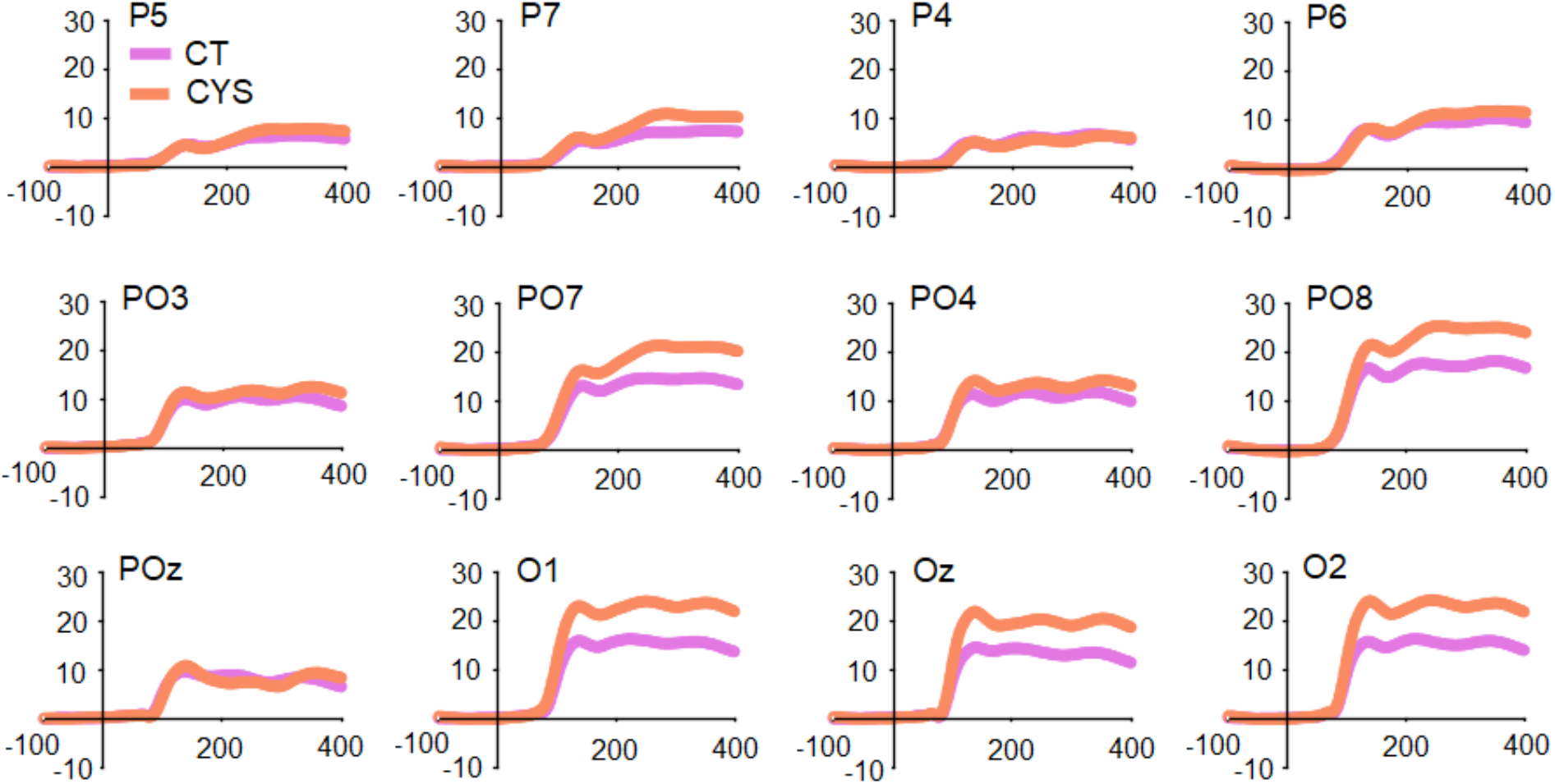
Averaged ERPs per group over parietal, parietal-occipital, and occipital channels.

## Notes

### Competing Interest Statement

The authors have declared no competing interest.

